# Prediction of the functional impact of missense variants in BRCA1 and BRCA2 with BRCA-ML

**DOI:** 10.1101/792754

**Authors:** Steven N. Hart, Eric C. Polley, Hermella Shimelis, Siddhartha Yadav, Fergus J. Couch

## Abstract

*In silico* predictions of missense variants is an important consideration when interpreting variants of uncertain significance (VUS) in the *BRCA1* and *BRCA2* genes. We trained and evaluated hundreds of machine learning algorithms based on results from validated functional assays to better predict missense variants in these genes as damaging or neutral. This new optimal “BRCA-ML” model yielded a substantially more accurate method than current algorithms for interpreting the functional impact of variants in these genes, making BRCA-ML a valuable addition to data sources for VUS classification.

## Introduction

Failure to accurately predict the effects of missense variants in *BRCA1* and *BRCA2* confound interpretation of gene sequencing studies and clinical care. Until recently, few missense variants had been functionally evaluated using validated assays, so interpretations of pathogenicity have relied on *in silico* predictions of functional effect in combination with family-based data. Many *in silico* prediction models are derived from supervised learning methods using variants in many different genes across the genome. The objective of supervised learning is to identify and weight a set of input features to correctly predict whether a variant is damaging, neutral, or somewhere in between.

Machine learning (ML) is a suite of computational algorithms that are able to parse data, learn higher dimensional representations of that data, and ultimately make a prediction using that data. A subclass of ML, known as supervised learning, involves utilizing a training dataset with known outcomes and learning a function to be able to evaluate new unknown outcome observations and make predictions of the outcome. Examples of machine learning include logistic regression algorithms and more complex ones like random forests, gradient boosting machines, and neural networks. Choosing the algorithms most suited to a particular task is an active area of research, since no single algorithm outperforms all others on every task.^1^ An efficient exploration of many different ML algorithms can be achieved through an automated machine learning (AutoML) approach.

A key limitation to the application of existing *in silico* models to assessment of variants in a specific gene is the reliance on known damaging variants in other genes. Such variants are likely to cause a number of different effects (e.g. alternative splicing, disruption of protein-protein interactions, altered protein folding, etc.) that may or may not be relevant for a given gene of interest. Gene specific models will likely outperform any general model, but only a few genes have been characterized to a degree that would be informative for single gene models. Two such exceptions are *BRCA1* and *BRCA2*. The landscape of functionally characterized variants in *BRCA1* has dramatically increased because of three major analyses. Starita et al.^2^, measured the impact of 1056 N-terminal variants on the homologous recombination DNA repair activity of *BRCA1*. Findlay et al.^3^, exploited the essentiality of BRCA1 for cell survival by used a saturating genome editing approach in HAP1 cells to evaluate nearly 4,000 SNVs (n=1837 distinct missense). Finally, Fernandes et al.^4^, reported on analysis of 354 distinct missense variants (n=79 in IARC classes 0 or 1 [benign] or 4,5 [pathogenic]) in the BRCT domain of BRCA1 using a validated transcriptional assay. Combined with results from a homology directed repair assay of 207 missense variants in the DNA binding domain of BRCA2,^5^ there are now sufficient numbers of variants to apply supervised learning methods to better predict damaging mutations in *BRCA1* and *BRCA2*.

## Methods

We employed the AutoML approach with the *R* (version 3.4.2) package *h2o.ai* (version: 3.16.0.2)^6^ to identify the optimal model for predicting the functional effect of missense variants in *BRCA1* and *BRCA2*. Variants were loaded in the following order: Hart^5^, Startia^2^, Fernandes^4^, and Findlay^3^; keeping only variants not observed in the previous studies. We also included new *BRCA2* functional data for 15 neutral (V2527A, G2544S, I2627V, M2634T, Y2658H, A2671S, I2675V, V2728A, P2767S, A2770T, A2770D, S2806L, I2822F, S3123R, Q2829R) and seven damaging mutations (F2562C, W2619G, K2657T, D2723N, L2753P, Y3006D, L3101R) (**Supplemental Table 1**). Since not all variants could unequivocally be assigned to a given class, we selected variants for inclusion if they satisfied the following criteria: “FUNC” (Neutral) or “LOF” (Damaging)^3^, HDR score <= 0.33 (Damaging) or >= 0.77^2^, or International Agency for Research on Cancer classes 0,1 (Neutral) and 4,5 (Damaging)^4^. Variants were excluded if they were not observed in known functional domains in *BRCA1* (BRCT: amino acids 1-109, RING: amino acids 1642-1855) or *BRCA2* (DNA-Binding: amino acids 2479-3192). This left 1902 variants (n=259 damaging) for *BRCA1* and 202 variants (n=74 damaging) for *BRCA2*.

For training each gene, 80% of variants were selected and trained to maximize the per class accuracy, with robustness assessed using 5-fold cross-validation. Input features were missense prediction models from dbNSFP (version 3.4)^7^, including SiftScore, Polyphen2HdivScore, Polyphen2HvarScore, LrtScore, MutationtasterScore, FathmmScore, ProveanScore, Vest3Score, MetasvmScore, MetalrScore, MCapScore, RevelScore, MutpredScore, CaddRaw, DannScore, FathmmMklCodingScore, GenocanyonScore, IntegratedFitconsScore, Gm12878FitconsScore, H1HescFitconsScore, HuvecFitconsScore, BayesDel, AlignGVGDPrior, EigenRaw, and EigenPcRaw. AlignGVGD^8^ and BayesDel^9^ were also added using the BioR framework.^10^ Optimal cutpoints for each of the individual input features (n=25) from dbNSFP, AlignGVGD, and BayesDel were determined using the same training data as used in AutoML so as to make a fair comparison.

For the test set evaluation, statistical measures of sensitivity, specificity were computed with the *caret* package^11^. The Matthews Correlation Coefficient (MCC) is used throughout as an optimal metric for gauging the performance of a binary classifier, as it represents a singular value that takes into consideration the proportion of each class. The values of MCC range from −1 to 1, where −1 represents the worst possible agreement and 1 representing perfect agreement. We also present traditional measures of performance for machine learning models such as receiver operating curves and precision-recall curves.

## Results

An iterative process was used to build hundreds of predictive models with different algorithms (Linear models, Gradient Boosting Machines, XGBoost, Neural Networks, Random Forests, Extremely Randomized Forests) and their associated hyperparameters. For *BRCA1*, 663 model/parameter combinations were tested with AutoML. The best performing model was a Gradient Boosting Machine with 48 trees of depth=8 and between 16-33 leaves. The mean MCC was 0.66 ± 0.049 s.d., corresponding to 89.5% sensitivity and 91.5% specificity. Similarly, for *BRCA2*, 76 model/parameter combinations were tested. The best performing model being an XGBoosted Machine with 50 trees with a mean MCC of 0.73 ± 0.057 s.d, sensitivity of 97.7% and specificity of 85.1%. Throughout the remainder of the manuscript, we will simply refer to these gene-specific models as BRCA-ML.

For the individual predictors (excluding BRCA-ML), the best model as determined by MCC for *BRCA1* was MutPredScore (MCC=0.399, sensitivity=94.5%, specificity=77.8%) and BayesDel (MCC=0.673, sensitivity=85.3%, specificity=83.0%) for *BRCA2*. More globally, we show the Receiver Operating and Precision-Recall Curves in **Figure 1**, which demonstrates better performance of BRCA-ML compared to other prediction models. In particular, the nigh number of false negative calls in BRCA1 many of the models yielded low area under the precision recall curves. BRCA-ML scores for every possible missense mutation caused by a single nucleotide variation are also given in **Supplemental Table 2**.

**Figure 1.**
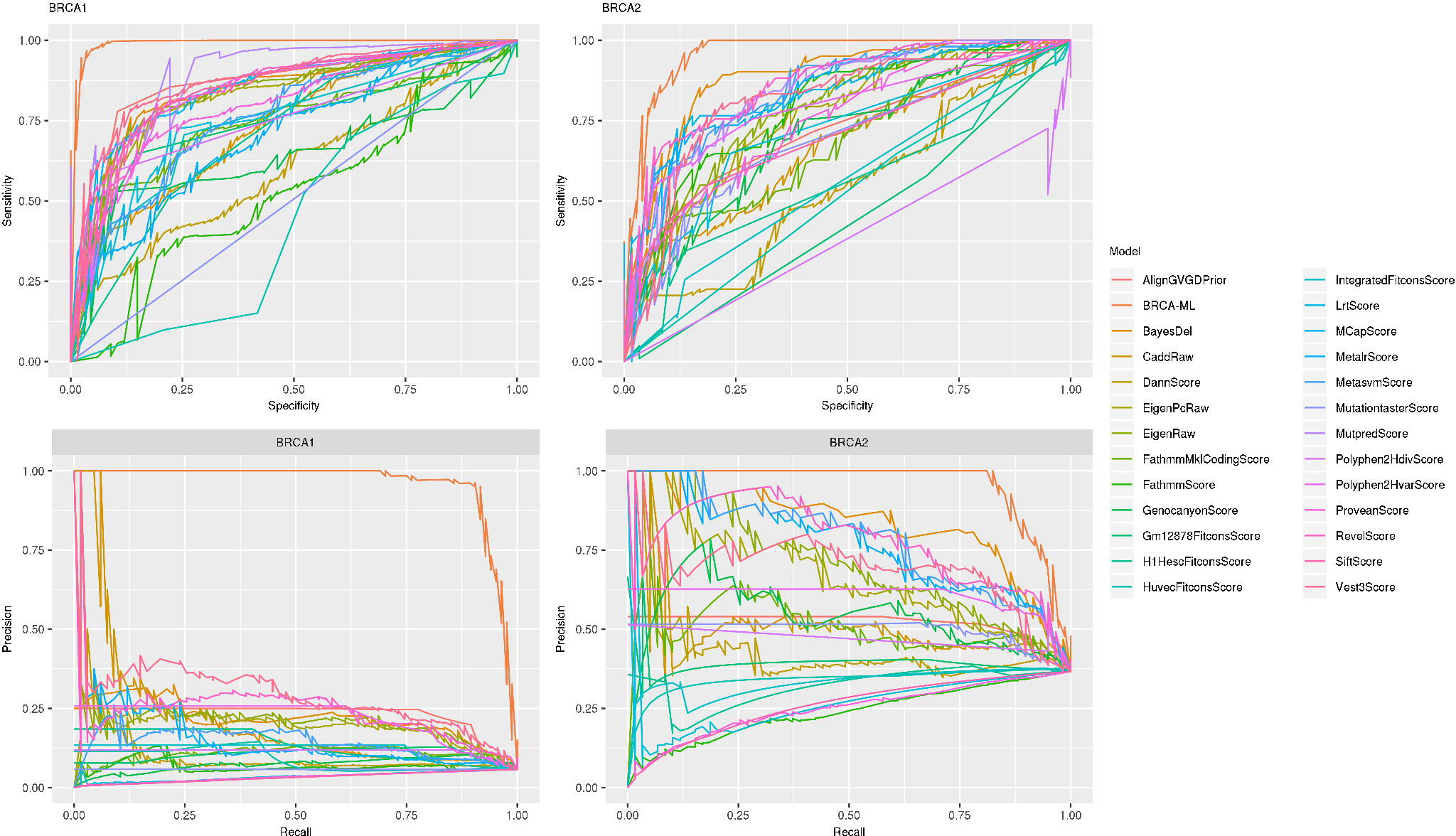
Receiver Operating Curves (top) and Precision-Recall Curves (bottom) for *BRCA1* (right) and *BRCA2* (left) for the hold out test set. The ideal location in the ROC curve is the top left corner, whereas the optimal position in the PR curve is the top right.

**Figure 2** shows the gene-level scores for every possible missense variant caused by a single nucleotide variant in *BRCA1* and *BRCA2* using BRCA-ML and BayesDel^9^, a commonly used and highly accurate predictor. While BayesDel is correctly assigning higher scores to known functional domains, the higher scores are not much more than predicted benign variants across the gene. However, in BRCA-ML, the signal to noise ratio is considerably higher between damaging and neutral variants. This evidence suggests that, unlike BayesDel, changing the threshold for damaging mutations will not significantly affect the number of predicted damaging mutations in BRCA-ML.

**Figure 2.**
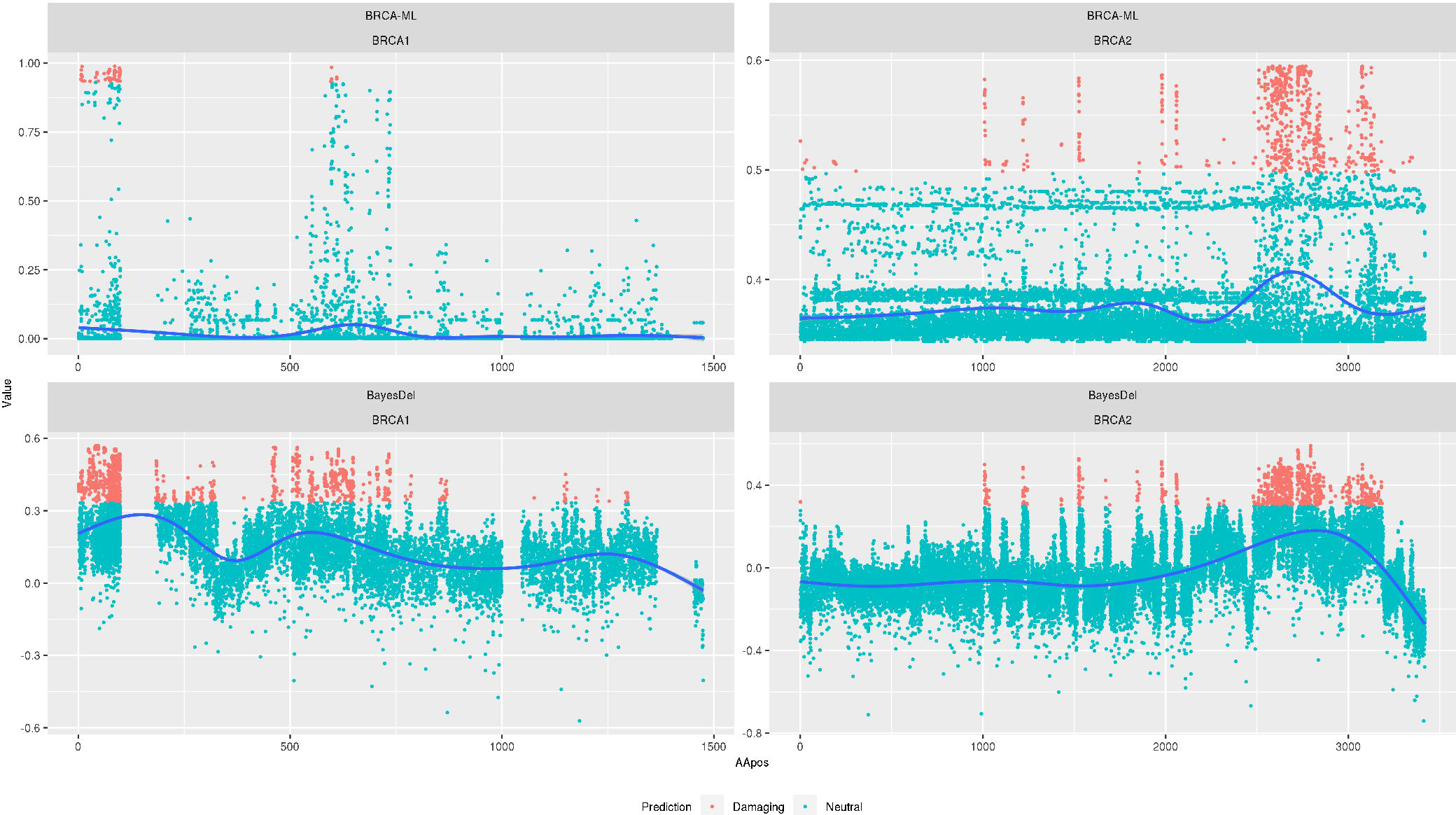
BayesDel (top) versus BRCA-ML (bottom) score distribution by gene. Distribution of missense prediction scores for *BRCA1* (left) and *BRCA2* (right) for each amino acid substitution. The blue line is a smoothed function to show where the density of scores is located.

## Discussion

We have shown that AutoML methods are efficient means to derive optimal machine learning models for predicting damaging missense mutations in *BRCA1* and *BRCA2*. The final models derived for each gene, which we collectively term BRCA-ML, show marked improvements in MCC and other metrics with respect to individual missense prediction algorithms.

Even in the age of large scale mutational scanning techniques like those from Findlay^3^ and Starita^2,3^, *in silico* mutation analysis will likely continue to be relevant. While the number of variants functionally tested is impressive for both studies (1,056 and 3,893, respectively), there are over 12,520 and 22,772 possible single nucleotide variants in *BRCA1* and *BRCA2*. Therefore, it could be several years before the technology exists to scale to all possible variants, hence a short term need for computational predictions.

It should be noted that there remain several limitations for these models. First, there are a limited number of known damaging mutations in *BRCA1* and *BRCA2* from which to build a model. The lack of damaging mutations limits the model ability to capture the complete variability of input data. Second, the training data are limited to characterized mutations in regions of the proteins known to be associated with impaired DNA damage repair. For example, the only missense variants in BRCA2 that are associated with disease are in the DNA binding domain. It is not known if variants in other domains that we or others predict to cause damaging missense mutations are able to inhibit DNA repair. Third, it is possible that there is some overfitting of the model due to the inherent biases in the 25 input features from dbNSFP. However, by keeping the test set isolated from the training data, this influence should be minimal. More known mutations in these genes will be necessary to quantify the amount of overfitting.

The data presented in this paper show that highly accurate prediction of missense variants in *BRCA1* and *BRCA2* are not only possible but simple to access (see **Supplemental Table 2** for all possible SNVs in both genes). This improved performance in BRCA-ML should provide higher quality evidence to genetic counselors and researchers for interpreting deleteriousness of missense variants.

## Code

All data and code are available at https://github.com/Steven-N-Hart/BRCA-ML

